# *U2AF1 S34F* enhances tumorigenic potential of lung cells by exhibiting synergy with *KRAS* mutation and altering response to environmental stress

**DOI:** 10.1101/2024.09.11.612492

**Authors:** Cindy E Liang, Eva Hrabeta-Robinson, Selam Mehreteab, Amit Behera, Colette Felton, Isobel J. Fetter, Cameron M. Soulette, Alexis M. Thornton, Shaheen S. Sikandar, Angela N. Brooks

**Author notes:** Co-first author (Cindy E Liang, Eva Hrabeta-Robinson).

## Abstract

Although *U2AF1^S34F^* is a recurrent splicing factor mutation in lung adenocarcinoma (ADC), *U2AF1^S34F^* alone is insufficient for producing tumors in previous models. Because lung ADCs with *U2AF1^S34F^* frequently have co-occurring *KRAS* mutations and smoking histories, we hypothesized that tumor-forming potential arises from *U2AF1^S34F^* interacting with oncogenic *KRAS* and environmental stress. To elucidate the effect of *U2AF1^S34F^* co-occurring with a second mutation, we generated human bronchial epithelial cells (HBEC3kt) with co-occurring *U2AF1^S34F^* and *KRAS^G12V^*. Transcriptome analysis revealed that co-occurring *U2AF1^S34F^* and *KRAS^G12V^* differentially impacts inflammatory, cell cycle, and KRAS pathways. Subsequent phenotyping found associated suppressed cytokine production, increased proliferation, anchorage-independent growth, and tumors in mouse xenografts. Additionally, HBEC3kts harboring *U2AF1^S34F^* alter splicing in stress granule protein genes, show increased stress granule formation, and have increased viability in cigarette smoke concentrate. Our results suggest that *U2AF1^S34F^* may potentiate transformation by granting precancerous cells survival advantage in environmental stress, permitting accumulation of additional mutations, like in *KRAS,* which synergize with *U2AF1^S34F^* to transform the cell.

## INTRODUCTION

Splicing factor mutations are prevalent in cancer and lead to global dysregulation of RNA splicing in protein-coding genes^1–10^. Work still remains to fully characterize the functional consequences of dysregulated splicing. *U2AF1* is among the most significantly mutated genes in lung ADC and codes for a splicing factor that is a subunit of the U2 Auxiliary Factor complex^11,12^. Of the *U2AF1* mutations in lung ADC, *U2AF1^S34F^* occurs the most frequently^2^.

Wild-type U2AF1 (U2AF1^WT^) facilitates spliceosome assembly by recognizing and binding to the 3’ splice site^13^. U2AF1 can also directly bind to mRNA to repress protein translation^14^. In *U2AF1^S34F^* cells, the amino acid substitution in the second zinc finger alters 3’ splice site choice^14–16^ and creates isoforms which result in abnormal gene expression^17^. Other impacts of this mutation include altered binding to mRNA leading to translational dysregulation, increased R loop formation, and reduced nonsens-mediated decay (NMD) activity^14,18,19^. Multiple functional consequences of this mutation have been reported, including increased survival advantage in cells exposed to ionizing radiation, altered inflammatory cytokine secretion, and increased stress granule production^14,20^. Despite these phenotypes, *U2AF1^S34F^* by itself is insufficient for human lung cell lines (HBEC3kts) to form tumors in mouse xenograft experiments^15^.

Confoundingly, *U2AF1^S34F^* is associated with poorer prognosis in cancer patients, but the mutation decreases proliferation in cancer cell lines^21^. In lung ADC, *U2AF1^S34F^*has been found to be a basal mutation^16,22^, indicating that it may potentiate the accumulation of further genetic perturbations under certain conditions or environments. Additionally, it has been observed to co-occur with known cancer-driver mutations like those in *KRAS* or *ROS1* fusions^11,16,22,23^. As such, understanding the impact of mutant *U2AF1* on oncogenesis in the context of different environments or with co-occurring mutations may pave the way for earlier diagnostic and treatment options for patients.

Although it has been hypothesized that *U2AF1^S34F^* confers tumorigenic potential independent of that conferred by the driver mutations with which it co-occurs^11^, work still remains to fully understand U2AF1’s functional significance. Mutations in *KRAS* associated with lung cancer, alone, are also insufficient to independently cause *in vivo* transformation in HBEC3kts^24^. *KRAS* mutations have recently been reported to alter splicing through downregulating splicing factor phosphorylation^25^. The cooperativity of splicing perturbations caused by co-occurring *U2AF1^S34F^* and *KRAS^G12V^* in a pre-cancerous model has yet to be studied.

Here, we introduced *KRAS^G12V^* to HBEC3kt lines with *U2AF1^S34F^*. We pair short-read mRNA sequencing with *in vivo* and *in vitro* assays to assess the impact of transcriptome alterations caused by co-occurring *U2AF1^S34F^* and *KRAS^G12V^* on preneoplastic potential and compare these perturbations to those caused by *U2AF1^S34F^*or *KRAS^G12V^* alone. Our results reveal synergistic effects of co-occurring *U2AF1^S34F^* and *KRAS^G12V^* in gene expression and splicing, which translate to enhanced oncogenic potential. Additionally, we discovered increased splicing in stress granule protein genes conferred by *U2AF1^S34F^* alone, which is associated with enhanced resistance to environmental stress. We propose a model for lung cancer oncogenesis in which *U2AF1^S34F^* enhances preneoplastic potential by allowing cells to survive stress and synergize with the transcriptomic effects of subsequent accumulated mutations.

## RESULTS

### *KRAS^G12V^* suppresses the effect of *U2AF1^S34F^* on the transcriptome while altering gene expression in oncogenic pathways

To understand the role of *U2AF1^S34F^* in cancer, we analyzed available data from lung ADC primary samples with *U2AF1* mutations to identify co-occurring mutations in known lung ADC driver genes. We find that *U2AF1^S34F^* significantly co-occurs with *KRAS* mutations (Figure 1A, Table S1), with *KRAS* mutations at the *G12* locus being the most common. From this analysis, we identified *KRAS^G12X^*as a candidate mutation to study in the context of *U2AF1^S34F^*.

**Figure 1:**
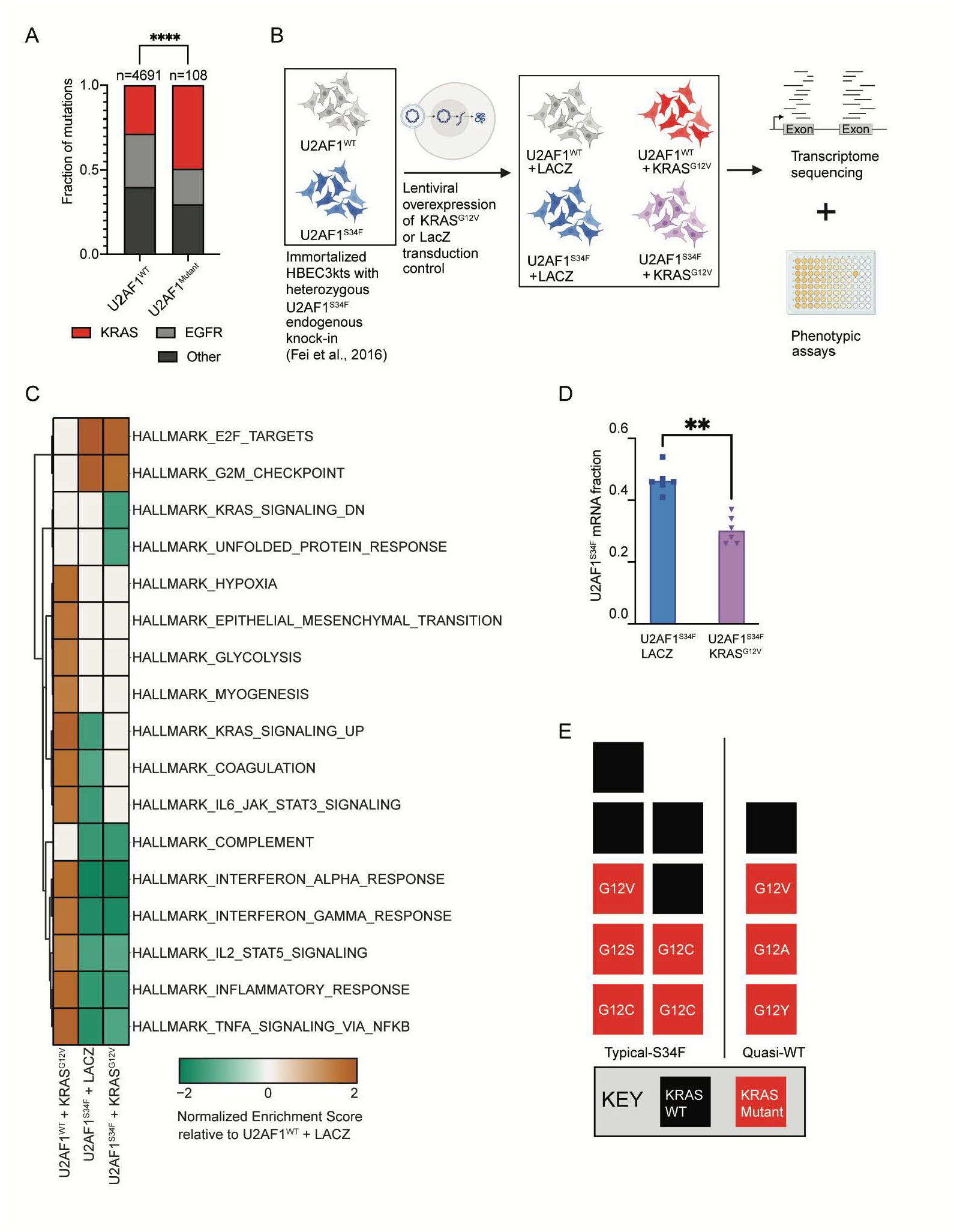
*KRAS^G12V^*suppresses the effect of *U2AF1^S34F^* on the transcriptome while altering gene expression in oncogenic pathways. **(A)** Distribution of *KRAS*, *EGFR*, and other mutations in lung ADC patients with and without mutations in *U2AF1*. (**B)** Experimental pipeline for study. Illumina RNA sequencing was performed on HBEC3kt lines with *U2AF1^S34F^* alone, *KRAS^G12V^*alone, co-occurring *U2AF1^S34F^* and *KRAS^G12V^*, and a wild-type control. Phenotypic assays for oncogenic phenotypes were also performed. (**C)** Heatmap of gene enrichment scores for gene sets differentially expressed between each genotype and the wild-type control. (**D)** *U2AF1^S34F^* mRNA fraction in HBEC3kt lines with differing mutational backgrounds. Bars represent mean *U2AF1^S34F^* mRNA fraction. (**E)** Distribution of *KRAS* mutations in lung ADC patients observed to display quasi-WT or typical-S34F expression patterns. Each box represents a single patient. ** P ≤ 0.01, **** P ≤ 0.0001. See also Figures S1-S2, Table S1, Table S2.

We obtained two parental isogenic HBEC3kt clones that were either wild-type or mutant for *U2AF1*^15^: one cell line was homozygous *U2AF1^WT^*, and the other cell line was heterozygous for *U2AF1^S34F^* at its endogenous locus (Figure S1 A). Homozygous *U2AF1^S34F^* mutation is lethal, so was not a consideraton^26^. We obtained a *KRAS^G12V^*pLenti6/V5 plasmid^24^ which has been used previously to identify genetic perturbations required to accomplish *in vivo* transformation of HBEC3kts, to study the impact of co-occurring *U2AF1^S34F^*and *KRAS^G12V^* mutations on preneoplastic potential. We exogenously overexpressed *KRAS^G12V^* in *U2AF1^WT^* and *U2AF1^S34F^* HBEC3kt cells using this construct. As a transduction control, we also introduced *LacZ* using the same plasmid backbone. A total of 4 cell lines, in biological triplicate, were generated per parental HBEC3kt clone: (1) *U2AF1^WT^* + *LacZ* (2) *U2AF1^S34F^*+ *LacZ,* (3) *U2AF1^WT^* + *KRAS^G12V^*, (4) *U2AF1^S34F^* + *KRAS^G12V^* resulting in sextuplicate cell lines per genotype. The presence of *U2AF1^S34F^* and *KRAS^G12V^* was validated through transcript expression and immunoblot (Figures S1 B-F).

Using these four cell lines, we performed short-read RNA sequencing and cell-based assays to understand how *U2AF1^S34F^* and *KRAS^G12V^* co-occurrence alters the transcriptome and biology of HBEC3kt (Figure 1B). We performed differential gene expression and gene set enrichment analysis (GSEA) on RNA-seq data (Figure 1C, Table S2). Gene sets uniquely downregulated in *U2AF1^S34F^* + *KRAS^G12V^*HBEC3kts were the KRAS Signaling Down gene set and the Unfolded protein Response gene set. The KRAS Signaling Down set corresponds to genes downregulated when KRAS is active, while the Unfolded Protein Response (UPR) set corresponds to genes up-regulated during an endoplasmic reticulum-related stress response^27^. In contrast, the *U2AF1^WT^* + *KRAS^G12V^* cell line had a positive enrichment for the KRAS Signaling Up gene set, corresponding to genes upregulated when KRAS is active, and no significant enrichment in the KRAS Signaling Down gene set. From these results, we infer that *U2AF1^S34^*^F^ in the presence of oncogenic KRAS alters KRAS and stress signaling pathways.

We also observed that, individually, *U2AF1^S34F^* and *KRAS^G12V^* produced opposite enrichment patterns to each other in KRAS signaling, coagulation, and inflammatory pathway gene sets (KRAS Signaling Up, Coagulation, IL6 JAK STAT3 Signaling, Inflammatory Response, TNFA Signaling Via NFKB, Interferon Alpha Response, Interferon Gamma Response, IL2 STAT5 Signaling). In some gene sets (Complement, IL2 STAT5 Signaling, Inflammatory Response, TNFA Signaling Via NFKB, Interferon Alpha Response, Interferon Gamma Response, E2F Targets, and G2M Checkpoint), the *U2AF1^S34F^*enrichment pattern persisted in the *U2AF1^S34F^ + KRAS^G12V^*cell line. In others (KRAS Signaling Up, Coagulation, IL6 JAK STAT3 Signaling), the opposite enrichment patterns caused by either mutation alone appeared to counter each other in the *U2AF1^S34F^ + KRAS^G12V^*line, producing nonsignificant enrichment in the gene sets. This suggests that *KRAS^G12V^*suppresses a subset of *U2AF1^S34F^*-specific gene expression signatures in the *U2AF1^S34F^*+ *KRAS^G12V^* cell line. To understand this observation in the context of allele expression patterns, we quantified the ratio of *U2AF1^S34F^*mRNA in cells with and without *KRAS^G12V^* using our short-read RNA-seq data. We found that the *U2AF1^S34F^* mRNA allele fraction in *U2AF1^S34F^* + *KRAS^G12V^* cells is reduced relative to *U2AF1^S34F^* alone (Figure 1D, Table S1). It has been previously reported that a subset of *U2AF1^S34F^* lung ADC primary samples have “quasi-WT” status, which represents tumors with low S34F:WT mRNA ratios, but unchanged absolute *U2AF1^S34F^* or total *U2AF1* mRNA levels^15^. These quasi-WT S34F:WT mRNA ratios range from 0.27-0.31 with concomitant *U2AF1* wild-type splicing patterns. This suggests that the presence of *KRAS^G12V^* may suppress *U2AF1^S34F^* allele expression to the levels seen in “quasi-WT” patient samples.

We then examined the mutational status of typical-S34F and quasi-WT samples from The Cancer Genome Atlas (TCGA) lung ADC cohort studied by Fei et al.^15^. Consistent with our hypothesis that mutant *KRAS* suppresses the *U2AF1^S34F^* allele expression signature, we found that quasi-WT patient samples carry a higher proportion of *KRAS* mutations (3/4 samples) than typical-S34F samples (5/9) (Figure 1E), although the difference is not statistically significant, likely due to low sample sizes. Together, these results support the hypothesis that *KRAS^G12V^*suppresses the *U2AF1^S34F^* transcriptomic signature by reducing the relative mutant allele expression.

### *U2AF1^S34F^* increases splicing in stress response genes

Although *U2AF1^S34F^* and oncogenic *KRAS* are both known to alter splicing^2,3,14–17,25,28–30^, the effect of co-occurring *U2AF1^S34F^* and *KRAS^G12V^* on differential splicing is unknown. First, we examined the mutations’ global effects on splicing.

We detected and quantified splicing events from short-read data using JuncBASE and compared our mutant cells to *U2AF1^WT^ + LacZ* ^32^. We found that, *U2AF1^S34F^ + LacZ* HBEC3kts exhibited the most differential splicing (Figure 2A). We found no significantly differentially spliced genes associated with expression of KRAS^G12V^, alone, which may partially be attributed to the U2AF1^WT^ + KRAS^G12V^ clones exhibiting the highest transcriptome variation among replicates (Figure S2A-B). *U2AF1^S34F^ + KRAS^G12V^* HBEC3kts had attenuated effects on splicing compared to U2AF1S34F + LacZ (Figure 2A). Consistent with the effect that *KRAS^G12V^* has on allelic expression of *U2AF1^S34F^* (Table S1), these results suggest that *KRAS^G12V^* suppresses the splicing changes mediated by *U2AF1^S34F^*.

**Figure 2:**
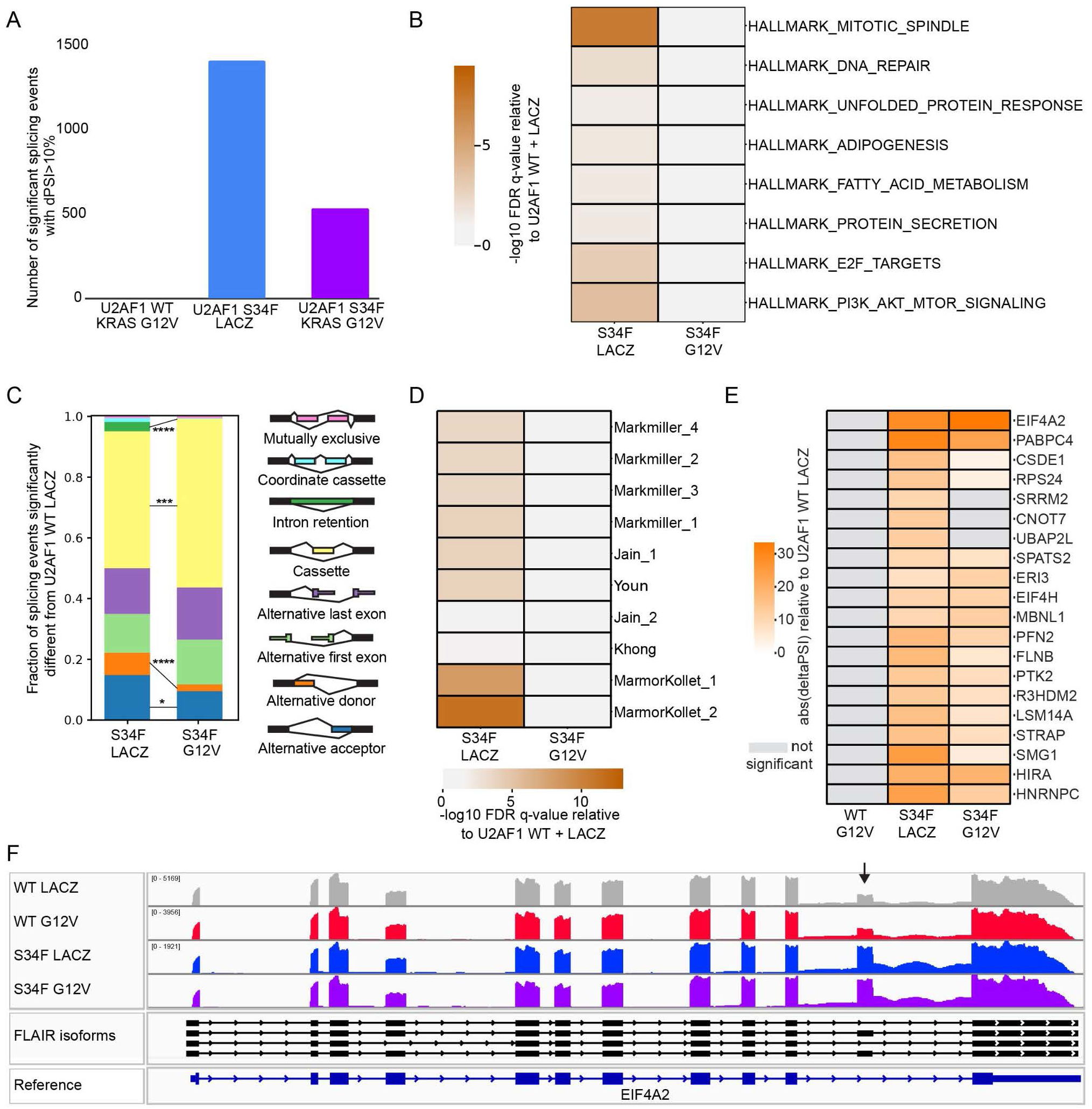
*U2AF1^S34F^*increases splicing in stress response genes. **(A)** Bar plot of number of differentially spliced genes (adjusted p < 0.25, delta PSI > 10%) compared to the wild-type control. (**B)** Heatmap of Hallmark gene set enrichment scores (unranked hypergeometric test) for differentially spliced genes (adjusted p < 0.05, delta PSI > 10%). **(C)** Stacked bar plot of event types of differentially spliced genes (adjusted p < 0.25, delta PSI > 10%). * P ≤ 0.05, ** P ≤ 0.01, *** P ≤ 0.001, **** P ≤ 0.0001 (**D)** Heatmap of stress granule gene set enrichment scores for differentially spliced genes (adjusted p < 0.05, delta PSI > 10%). **(E)** Heatmap of delta PSI values relative to U2AF1 WT LACZ for genes in significant stress granule gene sets that have also been shown to bind U2AF1. Non-significant changes are colored grey. **(F)** IGV snapshot of RNA-seq coverage of EIF4A2 in one representative replicate of each condition. Full-length isoforms assembled from previous long-read cDNA sequencing in these cell lines are shown (FLAIR isoforms, Soulette et al. 2023) Black arrow points to the poison exon.

To examine potential biological pathways impacted by the differentially spliced genes, we performed unranked GSEA on differentially spliced genes (Fig 2B). Similarly to the differential expression results, we see splicing impact cell cycle (Mitotic spindle, E2F targets) and inflammation pathways (PI3K, AKT, MTOR Signaling). However, we see in the gene expression that cell cycle pathways have increased expression in *U2AF1^S34F^* cells, while inflammatory pathways have decreased expression in *U2AF1^S34F^* cells. This indicates that *U2AF1^S34F^*-driven splicing changes can have differential impacts on gene expression.

We next compared the categories of altered splicing events that were significantly different (Fishers exact test p-value < 0.05) between *U2AF1^S34F^*+ *LacZ* and *U2AF1^S34F^* + *KRAS^G12V^*. JuncBASE categorizes splicing events in eight different categories: cassette exon, mutually exclusive exon, coordinate cassette exons, alternative 5’ splice site, alternative 3’ splice site, alternative first exon, alternative last exon, and retained intron (Figure 2C, Table S1). We found that co-occurring *U2AF1^S34F^*and *KRAS^G12V^* mutations cause predominantly cassette exon events, similar to *U2AF1^S34F^* alone, a splicing event type characteristic of *U2AF1^S34F^* ^2,14,15,17,29,30^ . This persistence of *U2AF1^S34F^*-specific splicing event was consistent with our gene expression results (Figure 1C), which demonstrated that certain *U2AF1^S34F^*-specific enrichment patterns persisted in the *U2AF1^S34F^ + KRAS^G12V^* line. In contrast, the intron retention, alternative donor, and alternative acceptor events significantly decreased in *U2AF1^S34F^ + KRAS^G12V^* HBEC3kts relative to *U2AF1^S34F^ + LacZ* cells. Overall, similar to gene expression enrichment patterns, we hypothesize that the effects of co-occurring *U2AF1^S34F^* and *KRAS^G12V^*mutations on splicing antagonize with each other to create intermediate splicing event proportions.

*U2AF1^S34F^* has also been found to confer altered splicing and binding to stress granule gene sets in a leukemia cell line^20^. Stress granules are RNA-protein condensates that may help cells survive stress and can help cancer cells resist chemotherapy^18^. In the leukemia line, altered splicing in stress granule genes was associated with enhanced viability under oxidative stress^20^. To test the hypothesis that *U2AF1^S34F^* alters stress response through aberrant splicing in our cell lines, we performed GSEA to analyze the enrichment of significantly spliced genes in genes coding for stress granule proteins (Fig 2D). We observed significant enrichment of stress granule gene sets in the *U2AF1^S34^*^F^ + *LacZ* cells, but not with the addition of *KRAS^G12V^*. We next investigated which stress granule genes in the significant gene sets were differentially spliced, restricting our search to only genes that are known to interact with U2AF1^14^. We found that many stress granule genes were significantly alternatively spliced in both *U2AF1^S34F^* + *LacZ* and *U2AF1^S34F^* + *KRAS^G12V^* cells, although fewer with the addition of *KRAS^G12V^*. One of the top alternatively spliced genes was *EIF4A2*, which contains a poison exon that would be predicted to be degraded by NMD and not produce protein. We found that the cassette poison exon of *EIF4A2* had significantly increased inclusion in both *U2AF1^S34F^* + *LacZ* and *U2AF1^S34F^*+ *KRAS^G12V^* cells, with concomitant decrease in total gene expression, as expected for a target of NMD.

### Altered splicing in stress granule genes in *U2AF1^S34F^* HBEC3kts is associated with enhanced stress response

Stress granules are often induced by an agent of oxidative stress^20,31^. Oxidative stress is relevant to cancer formation because it can produce mutations by causing DNA damage^32^. In lung ADC patients, a common source of oxidative stress is exposure to cigarette smoke. To understand the connection between oxidative stress response and lung ADC, we next analyzed splicing alterations in primary sample data from TCGA. We found greater numbers of splicing alterations in lung ADC samples from patients with smoking histories than in never-smokers (Figure 3A). Although we observed a trend of increased *U2AF1* mutations in samples from patients with smoking histories, high splicing alterations were present in *U2AF1^WT^* samples in this group as well.

**Figure 3:**
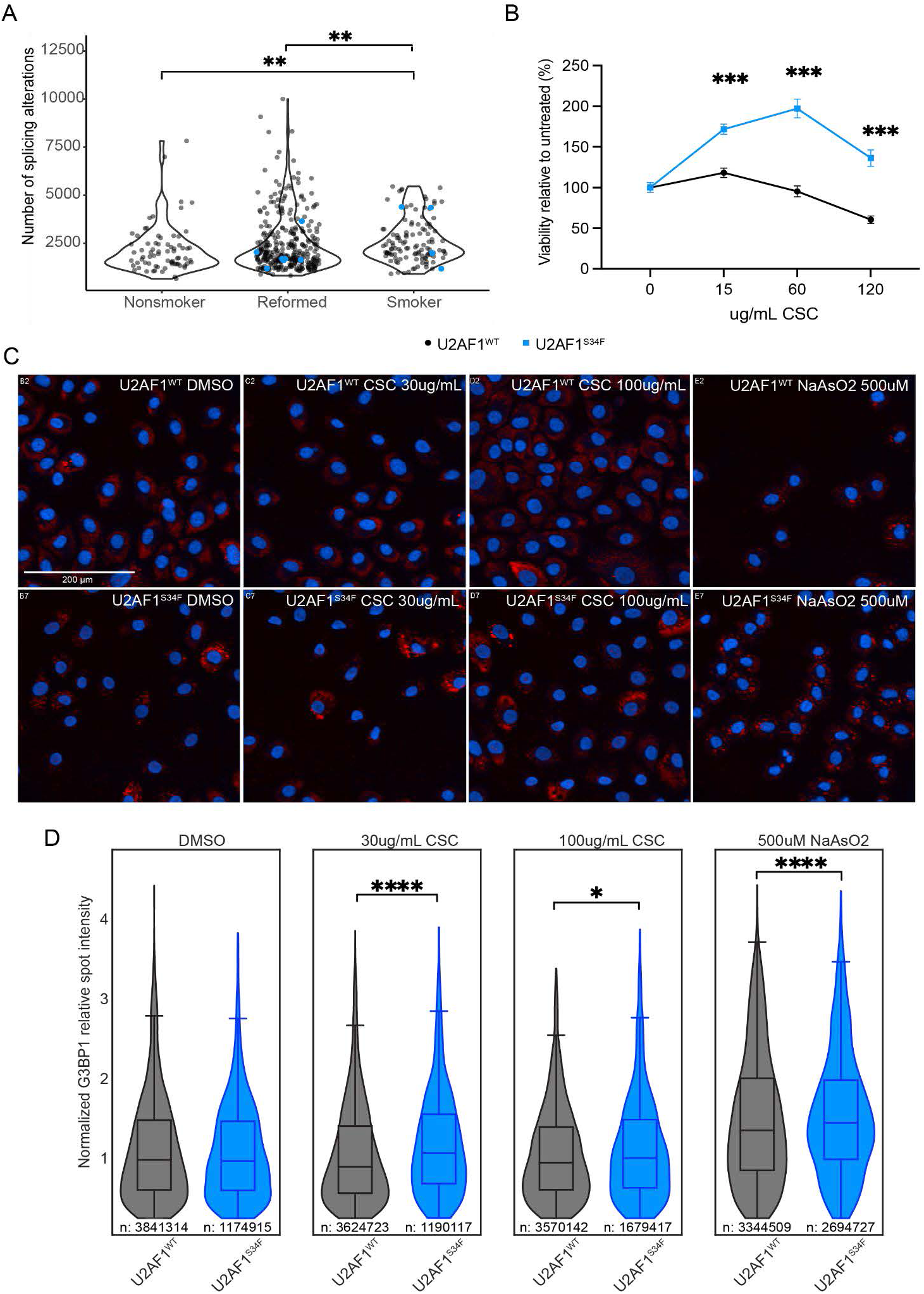
Altered splicing in stress granule genes in *U2AF1^S34F^* HBEC3kts is associated with enhanced stress response. **(A)** Splicing alteration distribution in lung ADC patients with smoking histories. Blue dots represent patients with *U2AF1* mutations. (**B)** Viability of HBECs with and without *U2AF1^S34F^* in cigarette smoke concentrate (CSC). Concentrations are in ug/mL CSC. Points represent mean viability of each cell line, while error bars show SEM. **(C)** Representative fluorescent images of G3BP1-positive granules (red) for HBECs mutant and wild-type for *U2AF1* under different treatment conditions. Hoechst stains showing the nucleus are in blue. DMSO = dimethyl sulfoxide, NaAsO_2_ = sodium arsenite. **(D)** Quantification of G3BP1 fluorescent intensity in wild-type and *U2AF1^S34F^* cells under different treatment conditions. The middle line in the body of each boxplot represent medians of each group, box limits represent quartiles, and whiskers represent the range of the most extreme, non-outlier data points. The number of G3BP1-positive spots imaged per group is listed below each plot. G3BP1 intensity is normalized by dividing all intensity values by the median relative spot intensity of *U2AF1^WT^* in DMSO. ** P ≤ 0.01, *** P ≤ 0.001. See also Data S2, Table S1.

Human cells harboring U2*AF1^S34F^* have observed an increase in viability after exposing cells to oxidative stress like radiation and sodium arsenite (NaAsO_2_), compared to *U2AF1^WT^* cells^14,20^. We followed up on this line of inquiry by measuring how *U2AF1^S34F^*alone impacts viability in cigarette smoke concentrate (CSC). We treated *U2AF1^S34F^*or *U2AF1^WT^* HBEC3kts with CSC and measured viability after three days. *U2AF1^S34F^* HBEC3kts displayed higher viability than *U2AF1^WT^* HBEC3kts at all concentrations tested (Figure 3B, Table S1, Figure S2D).

Next, we tested the hypothesis that *U2AF1^S34F^* lung cells alters stress granule abundance as a mechanism of enhanced stress resistance. We treated *U2AF1^S34F^* and wild-type HBEC3kts with CSC and stained cells for G3BP1, a stress granule component, to measure the intensities of cytoplsamic G3BP1-positive granules. DMSO was used as a negative control and NaAsO_2_ was used as a positive control (Fig 3C). The relative intensity of G3BP1-positive granules to the background was measured and normalized to the median relative intensity of *U2AF1^WT^* HBEC3kts in DMSO (Fig 3D). We observed increased G3BP1 intensities in *U2AF1^S34F^* HBEC3kts at both concentrations of CSC tested, as well as in NaAsO_2_.

Taken together, our data suggest that *U2AF1^S34F^* alters splicing in genes participating in stress granule formation to enhance the cell’s ability to respond to stress, including cigarette smoke exposure, granting *U2AF1^S34F^*cells a selective advantage.

### Co-occurring *U2AF1^S34F^* and *KRAS^G12V^* mutations increase oncogenic potential and proliferation

We next sought to understand the functional consequences of transcriptomic changes when *KRAS^G12V^* co-occurs with *U2AF1^S34F^*. We first sought to explore gene sets with similar enrichment patterns in both *U2AF1^S34F^ + LacZ* and *U2AF1^S34F^* + *KRAS^G12V^* HBEC3kts, as they indicated *U2AF1^S34F^*-specific effects which persisted when *KRAS^G12V^* was present. One category that fit this criteria was the inflammatory pathway gene sets (Complement, IL2 STAT5 Signaling, IL6 JAK STAT3 Signaling, Inflammatory Response, TNFA Signaling Via NFKB, Interferon Alpha Response, and Interferon Gamma Response), where the presence of *U2AF1^S34F^* was associated with downregulation. Oncogenic Ras has been found to increase production of cytokines such as IL-6 in multiple cell types^33,34^. To probe how these pathways are altered in our *U2AF1^S34F^* + *KRAS^G12V^*cell line, we measured inflammatory cytokine production in our HBEC3kt cell lines. For most cytokines tested, we observe that *U2AF1^WT^* + *KRAS^G12V^* HBEC3kts secrete the highest levels of inflammatory cytokines (Figure 4A). Co-occurrence of *U2AF1^S34F^*with *KRAS^G12V^* suppresses the levels of secreted cytokines IL-1β, IL-6, IL-8, TNFα, GM-CSF, and IFNγ (Figure 4A, Table S1). High levels of IL-1β, TNFα, GM-CSF, and IFNγ have been shown to promote antitumor activity in animal models^35–38^. We hypothesized that *U2AF1^S34F^*creates a microenvironment conducive to tumor growth by bringing cytokine secretion down to an intermediate level in *KRAS^G12V^*-mutant cells. We note that our cytokine results are inconsistent with previous work done on *U2AF1^S34F^* HBEC3kts, which showed that *U2AF1^S34F^*increases the secretion of cytokines such as IL-8^14^. However, the clonal background of the cells used in the aforementioned study was not reported, and it is possible that different steady-state cytokine secretion levels may be present in HBEC3kts from different isogenic clones.

**Figure 4:**
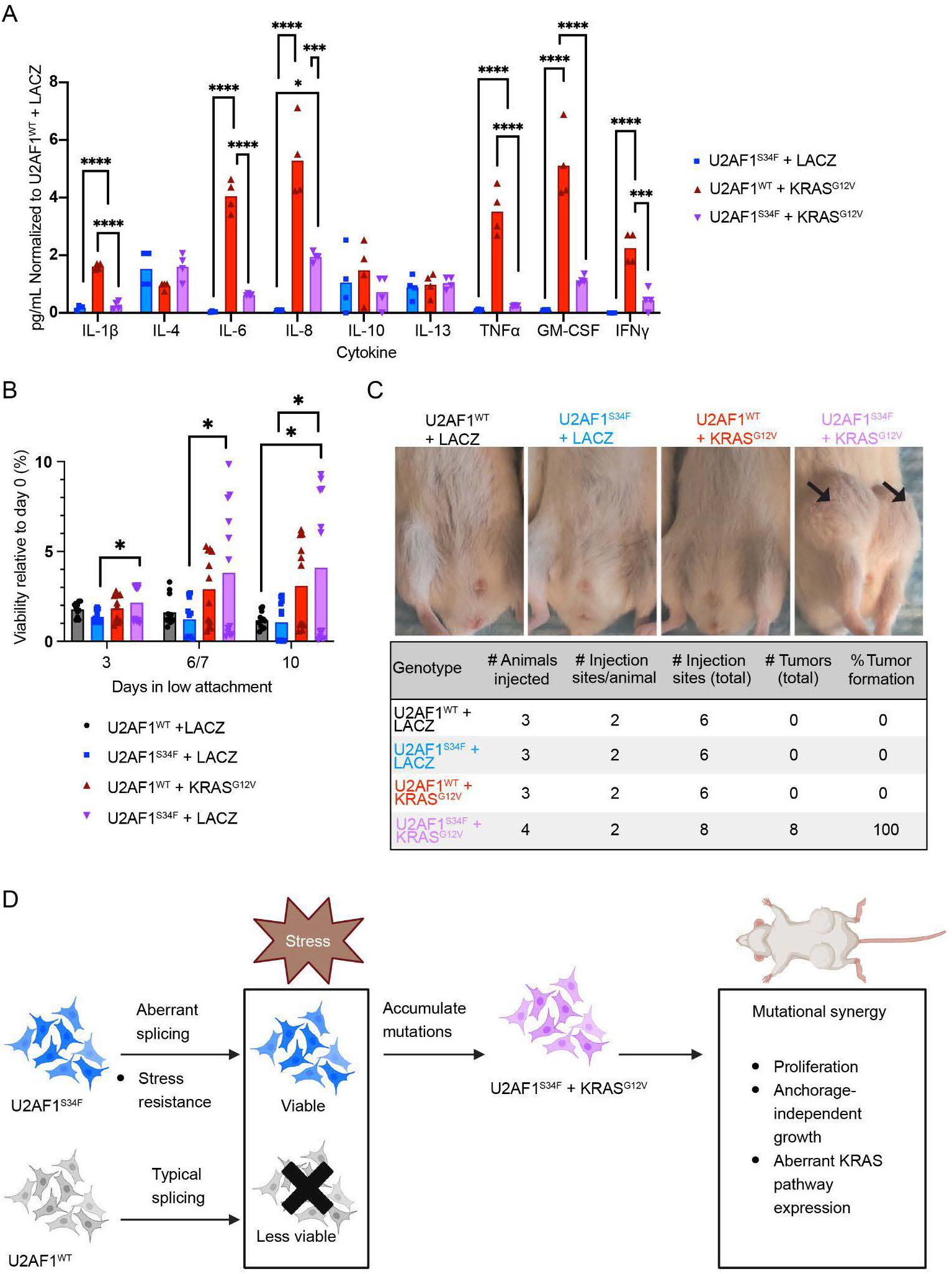
Co-occurring *U2AF1^S34F^* and *KRAS^G12V^* mutations increase oncogenic potential and proliferation. **(A)** Secreted cytokine measurements of each genotype, normalized to *U2AF1^WT^ + LACZ* levels. Bars represent mean normalized cytokine concentration. (**B)** Viability in low-attachment vessel for each HBEC3kt genotype. Relative viability is calculated by dividing the viability for each genotype at a certain time point, by the genotype’s viability at day 0. Bars represent mean viability. (**C)** Top, representative injection site images of mouse xenografts. Bottom, tumor formation quantification for each HBEC3kt genotype injected. **(D)** Working model for *U2AF1^S34F^*’s role in priming cells for oncogenic transformation. * P ≤ 0.05, ** P ≤ 0.01, *** P ≤ 0.001, **** P ≤ 0.0001. See also Figure S3B-E, Table S1, Table S3.

HBEC3kts with *U2AF1^S34F^* also exhibited enriched expression and splicing in gene sets related to cell cycle progression (E2F Targets, G2M checkpoint, and mitotic spindle). To understand how these gene expression differences translate to altered phenotype, we performed propidium iodide stainig and flow cytometry to measure the number of cells in each stage of the cell cycle (Fig S3). We observed small but significant changes in S phase between *U2AF1^WT^ + LacZ, U2AF1^S34F^ + LacZ*, and *U2AF1^wt^ + KRAS^G12V^* HBEC3kts.

There are known limitations in the ability of single-parameter DNA analysis to be able to distinguish between cells in very early or late S phase from cells in the G1 and G2 phases respectively^39^. Additionally, limitations exist in the ability of propidium iodide-based flow cytometry to distinguish between G2 and Mitotic phase cells^40^. To further understand putative cell cycle differences in HBEC3kts, we stained HBEC3kts with EdU and phospho histone-H3 (PHH3) to measure the proportion of cells undergoing S-phase and M-phase, respectively^40,41^ (Figures S4A-B, Table S3). Previous models with *U2AF1^S34F^* have found that *U2AF1^S34F^* by itself suppresses growth phenotypes such as proliferation and colony-forming potential^15,21^.

Consistent with these findings, we observed lower normalized EdU and PHH3 intensity in *U2AF1^S34F^* + *LacZ* HBEC3kts compared to *U2AF1^WT^* + *LacZ*. However, when *KRAS^G12V^* and *U2AF1^S34F^* co-occur, we observe increased proliferation compared to *U2AF1^S34F^* by itself.

Notably, M-phase staining in *U2AF1^S34F^* + *KRAS^G12V^* HBEC3kts was elevated to above *U2AF1^WT^* + *LacZ* levels (Figure S4B), indicating that *KRAS^G12V^* confers increased mitosis in *U2AF1^S34F^*-mutant cells. We note that the effect size of these changes are small but significant, and that other authors studying the *U2AF1^S34F^* mutation in different cell line contexts also report subtle changes in cell cycle^30^.

Mutant *KRAS*, including *KRAS^G12V^* is known to induce oncogenic phenotypes, such as anchorage-independent growth^41^. Due to the enhanced proliferation in *U2AF1^S34F^*+ *KRAS^G12V^* cells and the unique negative enrichment score in the KRAS Signaling Down gene set observed in this line, we hypothesized that co-occurring *U2AF1^S34F^* and *KRAS^G12V^* would alter anchorage-independent growth as well. Traditionally, this phenotype has been measured in HBECs by culturing cells in soft agar and quantifying the number of colonies formed^24^. However, a limitation of this assay is that it does not take into account metabolic activity as a metric for growth. We cultured HBEC3kts of the four genotypes on low attachment plates and measured growth over the course of 10 days using an ATP-based luminescent reagent^42^. *U2AF1^S34F^* + *KRAS^G12V^* HBEC3kts survived anchorage-independent growth conditions better than other genotypes over 10 days in low-attachment conditions (Figure 4B, Table S1).

We also assayed for other cancer hallmarks, such as the long-term ability to survive and proliferate into colonies, which is a marker of cancer stemness and can be assessed with a clonogenicity or colony-forming assay^43^. Interestingly, co-occurring *KRAS^G12V^*and *U2AF1^S34F^* suppressed colony-forming potential (Figure S4C, Table S1). These findings are consistent with previous work performed on *U2AF1^S34F^*-mutant cancer cell lines^21^. We also performed a wound-healing assay to assess the invasive potential of *U2AF1^S34F^* + *KRAS^G12V^*HBEC3kts (Figure S4D, Table S1) and observed that *U2AF1^S34F^* decreases invasive potential in *KRAS^G12V^* background. Our results indicate that enhanced proliferation conferred by *U2AF1^S34F^* + *KRAS^G12V^* may work in concert with pathways outside of stemness and invasion to confer oncogenic potential.

Finally, we sought to understand how *U2AF1^S34F^* and *KRAS^G12V^* co-occurrence in HBEC3kts impacts the ability of cells to form tumors, *in vivo*. Cells from the four genotypes were injected into NOD scid gamma (NSG) immunodeficient mice. We find that *U2AF1^S34F^*+ *KRAS^G12V^* HBEC3kts formed more tumors than HBEC3kts with either mutation alone (Figure 4C). This suggests that co-occurring *U2AF1^S34F^* and *KRAS^G12V^* mutations synergize to transform HBEC3kts cells *in vivo*. Importantly, no tumors were formed with *U2AF1^S34F^*, suggesting that *U2AF1^S34F^* alone is insufficient for *in vivo* transformation.

Together, our results lead us to a model of oncogenic transformation. *U2AF1^S34F^* has been reported to be a truncal mutation in lung cancer and MDS^16,22,44^. We propose that *U2AF1^S34F^*, when present in precancerous cells, allows cells to survive under oxidative stress. The surviving cells are more likely to persist and accumulate further mutations like *KRAS^G12V^*, which act synergistically with *U2AF1^S34F^* to alter splicing and gene expression, resulting in an increase in oncogenic potential (Figure 4D).

## DISCUSSION

Despite its status as a recurrent mutation, the role of *U2AF1^S34F^* in lung cancer has been difficult to understand since the mutation confers anti-proliferative and anti-invasive phenotypes when present alone in model systems. This aspect limits the ability of researchers to identify the functional role of *U2AF1^S34F^* in lung cancer and limits the use of *U2AF1^S34F^* as a prognostic marker for lung ADC. To gain a better understanding of this mutation, we examined the role of *U2AF1^S34F^*in early cancer formation in two directions: how *U2AF1^S34F^* may synergize with other cancer drivers like *KRAS^G12V^*, and how *U2AF1^S34F^*by itself can impact stress response in the cell.

Validation of pathways predicted to be differentially altered by gene set enrichment analysis revealed that the direction of enrichment in a gene set did not always correspond with the direction of pathway change in phenotype. For instance, although cell cycle gene sets were both positively enriched in *U2AF1*-mutant cell lines regardless of *KRAS* status, *U2AF1^S34F^* + *LacZ* cells exhibited reduced proliferation. In contrast, the co-occurrence of *U2AF1* and *KRAS* mutations increased proliferation. Other gene set enrichment patterns translated to more consistent phenotypes. For example, a negative enrichment in inflammatory gene sets translated to suppression of inflammatory cytokines for HBEC3kt lines.

We also examined splicing-level changes in the transcriptome caused by *U2AF1* and *KRAS* mutations. Interestingly, we found differences in gene set enrichment between differentially expressed and differentially spliced genes, highlighting the importance of using multiple kinds of RNA-seq analysis to assess the synergistic impact of mutations. For example, the P53 pathway was found to be significantly altered through splicing but not gene expression analysis, with an even stronger enrichment score in U2AF1^S34F^ + KRAS^G12V^ cells. We also detected *U2AF1^S34F^*-specific splicing event types, such as cassette exon event usage, that persisted in *U2AF1^S34F^ + KRAS^G12V^* cells, while *KRAS^G12V^* presence suppressed other *U2AF1^S34F^*-specific event types to an intermediate level.

When we followed up on our splicing analysis by quantifying cellular stress response, we found that *U2AF1^S34F^* confers enhanced viability under CSC treatment, which corresponds with an increase in G3BP1-positive granules. Our work leads us to a model in which *U2AF1^S34F^*confers a selective advantage that is dependent on the presence of environmental stress and that this advantage is mediated through altered splicing in stress granule genes. Resistant cells are then likelier to accumulate additional oncogenic drivers like *KRAS^G12V^*, which synergize with *U2AF1^S34F^* to increase oncogenic potential by altering inflammatory and anchorage-independent phenotypes.

Our findings of synergistic functional effects of *U2AF1* and *KRAS* mutation is even more intriguiging given a recent study that found that *KRAS^G12S^* mutation causes exon skipping in *KRAS* itself, which is then rescued by the presence of a *U2AF1* mutation ^45^. In our study, we find that the introduction of *KRAS^G12V^* suppresses the *U2AF1* mutant allele expression and subsequent effects on splicing. Future work to better understand the mechanism of this allele suppression phenotype will help to better understand the functional synergies of these two mutations.

More work is left to be done to understand the role of splicing in stress response. For instance, the persistence of cassette exon events in *U2AF1^S34F^ + KRAS^G12V^* cells highlights this category of splicing events as an interesting candidate for further functional study. Intron retention (IR) is another splicing event type linked to stress response in eukaryotes. In yeast, IR has been linked to fitness advantage in the presence of environmental stressors like starvation^46^. In mouse cells, IR has been linked to osmotic stress^47^. Although we did not find evidence of altered intron retention in our short-read analysis, previous work from our group utilizing long-read sequence analysis has shown *U2AF1^S34F^* increasing IR from long-read data^17^, leaving an interesting avenue to pursue as long-read sequencing technologies improve.

Overall, our study points to the importance of interrogating the function of cancer mutations in the context of other mutational contexts and environmental conditions to obtain a more complete understanding of their contributions to tumorigenesis.

## Lead contact

Please direct all manuscript questions to Dr. Angela Brooks

## Material availability

Cell lines generated from this study are available upon request.

## Data and code availability

Sequence data have been deposited at GEO under GSE267349. JuncBASE splicing quantification of cell lines and primary TCGA samples are deposited in Zenodo:https://zenodo.org/records/20514236. All code have been deposited to GitHub is publically available at https://github.com/cindyeliang/u2af1-kras/tree/main/scripts.

## EXPERIMENTAL MODEL DETAILS

### Mouse models

*NOD.Cg-Prkdc^scid^Il2rg^tm1Wjl^/SzJ* (NSG) (stock #005557) mice were purchased from The Jackson Laboratory and bred at the University of California Santa Cruz (UCSC). All mice used for this study were maintained at the UCSC Animal Facility in accordance with the guidelines set forth by UCSC and the Institutional Animal Care and Use Committee (Protocol number SIKAS2010).

## METHOD DETAILS

**Cell lines:** Host HBEC3kt cell lines homozygous for wildtype *U2AF1* and heterozygous with one copy of *U2AF1*^S34F^ at the endogenous locus were obtained as a gift from the laboratory of Harold Varmus (Cancer Biology Section, Cancer Genetics Branch, National Human Genome Research Institute, Bethesda, United States of America and Department of Medicine, Meyer Cancer Center, Weill Cornell Medicine, New York, United States Of America) and maintained as described by Fei et al.^15^. We received two isogenic clones each of the U2AF1^S34F^ and U2AF1^WT^ HBEC3kt lines, as described in their initial publication, and performed subsequent genetic manipulations on the clones separately^15^.

### KRASG12V and LACZ transduction

Clone 1 and clone 2 of HBEC3kt lines were used for lentiviral transduction and blasticidin selection to generate a stable expression of *KRAS*^G12V^ or *LACZ* using plasmids obtained as gifts from the laboratory of John D Minna (Hamon Center for Therapeutic Oncology Research, The University of Texas Southwestern Medical Center). Cells were selected and cultured as described in Sato et al.^25^. Cell lines generated were tested for mycoplasma (IDEXX). Biological replicates were cultured from each clone for each genotype (*U2AF1^WT^ + LACZ, U2AF1^S34F^ + LACZ, U2AF1^WT^ + KRAS^G12V^, U2AF1^WT^ + LACZ)*.

### RNA extraction of HBEC3kts

For RNA sequencing of HBEC3kt lines, cells were allowed to recover from cold storage in liquid nitrogen after seeding for one passage in a T-25 flask. Cells were passaged at 70-80% confluency to maintain log-phase growth into a 10cm plate. Once cells in the 10cm plates reached 70-80% confluency, they were washed twice in ice-cold DPBS and then collected in Tri-reagent for storage at -80°C until the bulk RNA was extracted using Direct-Zol RNA Miniprep Kit (Cat#R2050, Zymo Research).

### Short-read RNA-Seq of U2AF1^WT^ + LACZ, U2AF1^S34F^ + LACZ, U2AF1^WT^ + KRAS^G12V^, and U2AF1^S34F^ + KRAS^G12V^ HBEC3kts

For Illumina sequencing, n=3 10cm plates per HBEC3kt genotype of both clones, for a combined n=6 per genotype, were cultured for RNA extraction as described above. Concentrations of purified RNA in nuclease-free water were determined by Nanodrop-2000 Spectrophotometer and Qubit RNA BR Assay (ThermoFisher Scientific). RINe numbers ranging from 7.8-10 were determined by TapeStation 4150 RNA ScreenTape Analysis (Agilent Technologies) before sending RNA to UC Davis DNA Technologies and Expression Analysis Core Laboratory for poly-A strand specific library preparation to obtain 60 million paired end read pairs by NovaSeq S4 (PE150) sequencing.

### Mouse xenograft of HBEC3kts

Clone 1 and clone 2 HBEC3kt lines were cultured as previously described^15^. Cells were allowed to recover from cold storage in liquid nitrogen after seeding for one passage in a T-25 flask. Cells were passaged to a 10cm plate, then to a final 15cm plate, and allowed to grow to 80% confluency. At 80% confluency, the media in the 15cm plates were aspirated, and cells were washed twice with DPBS. To suspend cells for injection, cells were trypsinized with standard protocols^48^, and live cell counts were assessed by Trypan Blue staining. For each cell line, 9 million cells were resuspended in Keratinocyte SFM media containing 40% Matrigel and subcutaneously injected into the fourth abdominal fat pads on both sides of male NSG mice. 2-5 million cells were injected at each site in 100 uL media + Matrigel (5 million in 1st xenograft experiment, 2 million in 2nd). Mice were monitored every week for tumor growth. All mice were euthanized if tumor growth reached end point (1500 mm^3^), the tumors were ulcerated, or mice showed signs of distress. Tumor size was measured using digital calipers. A total of 5 female and 15 male mice were used.

### Viability assay

Clone 1 and clone 2 HBEC3kts of differing genotypes were seeded in a 96 well-plate in triplicate and grown in supplemented Keratinocyte Serum-Free Media (KSFM) (Cat# 17005042, Invitrogen). At multiple time points, (0, 4, and 6 days), cells were rinsed twice with DPBS, CellTiter-Glo (Cat# PRG7572, Promega) reagent was added, and cells were transferred to white opaque 96 well-plates for luminescence measurement. Luminescence at each timepoint was quantified using the VarioSkan platereader (Thermo Scientific) and normalized to the average relative luminescence units (RLU) of the 0 day timepoint.

### Western blot analysis

Clone 1 and clone 2 cell lines were cultured to 85% confluency in 10cm plates. After preparation of protein lysates in 1ml of RIPA buffer supplemented with protease inhibitor cocktail (Cat# 5892970001, Roche Molecular Systems, Inc, USA) proteins were denatured using standard denaturation techniques in beta mercaptoethanol laemmli buffer, and 15ug of denatured protein lysate was separated on a 4-15% Mini-Protean TGX Precast Protein Gel (Cat# 4561086, Bio-Rad Laboratories, Inc. USA). After transfer to 0.2 um PVDF membrane using TransBlot Turbo Transfer system (Cat# 1704272, BioRad Laboratories), membranes were incubated shaking at room temperature in 5% milk block in 1x PBST followed by incubation in KRAS^G12V^ primary antibody at 1:250 dilution (Cat# 14412, Cell Signaling Technologies) and B actin conjugated to HRP at 1:500 dilution (Cat# sc-47778 HRP, Santa Cruz Biotechnology) in milk block overnight on an orbital shaker at 4°C.The next day, blots were washed in PBST and incubated with secondary HRP-conjugated antibody (Cat# 7074, Cell Signaling Technologies) at 1:1000 dilution at room temperature for 1h. After washing in PBST, bands were detected using WesternSure PREMIUM Chemiluminescent substrate (Cat# 926-95000, Li-COR Biosciences) and visualized on a C-Digit Blot Scanner (Li-COR Biosciences).

### Secreted cytokine analysis

Growth triplicates of clone 2 cell lines were seeded in 6 well plates and cultured with standard protocols described above to 85% confluency. Conditioned media (3 mL) above the cells was collected and cell debris spun out at 3000 x g for 10 mins at 4°C and supernatent was stored in -80°C before sending to Eve Technologies (Calgary, Canada) for the Human High Sensitivity T-Cell Discovery Array 14-plex (HDHSTC14) assay, which measures cytokine concentrations using a bead-based ELISA. Data was plotted and significance was calculated with a Mann-Whitney test on GraphPad Prism.

### Cell cycle assays

Growth triplicates of clone 2 cells were seeded in a tissue culture-treated T-25 flask and allowed to reach a minimum of 50% confluency before collection of ∼1M cells by trypsinization. After washing cells 1x in DPBS, the cell pellets were fixed by vortexing with a dropwise addition of 2 mL of ice cold 70% ethanol. Fixed cells were stored in 4°C until ready for flow cytometry. To perform flow cytometry, fixed cells were transferred to capped polystyrene tubes and centrifuged at 2000 rpm for 5 min in an eppendorf 5810k centrifuge. The supernatant was removed and fixed cells were washed 1x in cold DPBS before final resuspension in 200uL of DPBS and 2 uL of RNAse A. 200uL of 50ug/mL of propidium iodide (PI) was added to the cell suspension. The final suspension was filtered through a 100 uM mesh before running through a LSRII flow cytometer. PI incorporation of the fixed cells were then measured using a 488nm laser.

### Proliferation immunofluorescent assays

Cell staining was performed at UCSC’s Chemical Screening Center, using the BioTek EL406 with peri/syringe/wash modules for automated washing and dispensing of reagents. Growth triplicates of clone 2 cells were cultured as previously described in optical-bottom black opaque 96 well-plates (Cat# 3904, Corning). The plate was taken to the Chemical Screening Center and incubated with EdU for 1 hour at 37°C and 5% CO2. Following EdU incorporation, cells were fixed with 5% formaldehyde (Cat# F79-500, Fisher) in basal media (Cat# PCS-300-030, ATCC) for 30 minutes at 37°C and 5% CO2. Cells were blocked with 2% BSA in PBST for 20-60min in the dark at room temperature.

Following blocking, click reagent (15ml 100mM Tris pH7.4, 0.6ml 100mM CuSO4, 155.5ul 200mg/ml Na Ascorbate, 15.5ul 10mg/ml Rhodamine-Azide) was added to the cells to visualize EdU incorporation and cells were incubated in the dark for 1h at room temperature. Following azide incorporation, cells were stained with Hoechst at 1:10000 dilution in 2% BSA in DPBS to visualize nuclei and incubated in the dark for 2h at room temperature. Cells were then incubated with a primary antibody for PHH3 (Ser10) Recombinant Rabbit Monoclonal Antibody (9H12L10) (Cat# 701258, Invitrogen) at 1:5000 dilution and mouse monoclonal anti α-Tubulin−FITC antibody (Cat# F2168-.2ML, Sigma-Aldrich) at 1:2500 dilution in 2% BSA in DPBS overnight at 4°C. Then, the plate was incubated in chicken anti-Rabbit IgG (H+L) Cross-Adsorbed Secondary Antibody, Alexa Fluor™ 647 secondary antibody (Cat# A-21443, Invitrogen) at 1:1000 dilution and Hoecsht at 1:20000 dilution in 2% BSA in DPBS for 2 hours in the dark at room temperature. The stained plate was then washed and stored in DPBS and 0.1% sodium azide at 4°C until imaging.

### EdU, Hoechst, and PHH3 imaging and analysis

Imaging and quantification of EdU, Hoechst, and PHH3 immunofluorescent signal was performed with the Perkin Elmer Opera Phenix Plus and Harmony bioinformatics software at UCSC’s Chemical Screening Center. Valid cells were identified as nonborder objects with the presence of nuclear Hoechst staining. The mean EdU and PHH3 intensities of each valid object were divided by the mean Hoechst intensities to normalize for cell density. N = 12 wells in a 96 well-plate were seeded per genotype and at least 36,000 objects were analyzed per genotype. The resulting values were plotted and statistical analysis was performed on Python.v3.7.7 and Jupyter notebook v6.3.0. Significance was calculated using a Kruskal-Wallis test with Dunn’s multiple comparisons test.

### Growth in Low Attachment (GILA)

Clone 1 and clone 2 cell lines were grown to 85% confluency on regular tissue culture-treated 6-well plates, harvested by trypsinization, filtered over a nylon 70um mesh and seeded in triplicate in KSFM media at 2500 cells per well in Ultra-low attachment 96 well plates (Cat# 3474, Corning) and time points were collected for viability assays over a 14-day period. An early time point was collected at the time of seeding and used for normalization. Viability was assayed using CellTiterGlo according to manufacturer’s instructions (Cat# G7570, Promega) and luminescence was measured on a VarioScan LUX plate reader (ThermoFisher). Significance was calculated using a Kruskal-Wallis test with Dunn’s multiple comparisons test using GraphPad Prism.

### Clonogenicity assay

Colony formation was assessed by seeding clone 2 cell lines in triplicate at 200 cells per 10cm plate and cultured under normal conditions, except that media was changed only twice over a 10 day period so as not to disturb colony formation. Cells were fixed in 100% methanol for 20 mins and stained in 0.5% Crystal Violet in 25% methanol for 5 mins before drying and photographing. Colonies of approximately 2mm or larger were counted in 4 separate quadrants of each plate. Significance was calculated using a Kruskal-Wallis test with Dunn’s multiple comparisons test using GraphPad Prism.

### Wound healing assay

Clone 2HBEC3kts were seeded in 6-well plates. A 200uL pipettor and filter tip was used to create the wound in a confluent monolayer of cells. The wound was imaged at 0 and 3 hours. The number of cells that had migrated into the wound between the two time points was counted. Significance was calculated using a Kruskal-Wallis test with Dunn’s multiple comparisons test using GraphPad Prism.

### Chemical stress treatment and stress granule immunofluorescent assays

Stress granule staining was performed using a protocol adapted from <u>kedersha and Anderson</u>. Clone 1 and clone 2 U2AF1WT + LACZ and U2AF1S34F + LACZ cells were cultured in optical-bottom black opaque 96 well-plates (Cat# 6055302, PerkinElmer). Cells were treated with 30ug/mL and 100ug/mL of CSC from Murty Pharmaceuticals (Cat# nc1560725) dissolved in KSFM to induce stress granule formation. As a positive control, cells were treated with 500 and 1000uM NaAsO2 (Cat# S7400, Sigma-Aldrich). Stock CSC is dissolved in DMSO, so an equal volume of DMSO as the volume of stock CSC added to the 100ug/mL CSC conditions was added to the negative control wells. Following treatment, the well-plate was incubated at 37°C for one hour. After this time period, wells were rinsed once with warm DPBS through gentle pipetting. To fix the cells, 36% formaldehyde (Cat# 47608-250ml-f, Sigma-Aldrich) was added and the plate was incubated at 37°C in a rotator for 30 minutes. Immediately following removal of formaldehyde, methanol (Cat# 106011, Millipore Sigma) pre-chilled at -20°C was added and the plate was incubated at room temperature for 10 minutes in a rotator to permeabilize the cells. Cells were blocked with 5% BSA in DPBS for one hour at room temperature. To visualize nuclei, cells were stained with Hoechst as described previously. To visualize stress granules, the plate was incubated in Rabbit anti G3BP1 at 1:300 dilution in 2% BSA in DPBS for one hour at room temperature. To visualize the cell body, cells were next stained with mouse monoclonal anti α-Tubulin−FITC antibody (Cat# F2168-.2ML, Sigma-Aldrich) at 1:2500 dilution in 2% BSA in DPBS for one hour at room temperature. The plate was then incubated in chicken anti-rabbit secondary Ab conjugated to Alexa Fluor 647 (Cat# A-21443, Invitrogen) at 1:1000 dilution and Rhodamine (TRITC) Bovine anti-goat IgG (Cat# 805-025-180, Jackson ImmunoResearch Laboratories Inc) at 1:100 dilution for 2 hours in the dark at room temperature. The stained plate was then washed and stored in DPBS and 0.1% sodium azide at 4°C until imaging.

### Stress granule imaging and analysis

Imaging and quantification of EdU, Hoechst, and PHH3 immunofluorescent signal was performed with the Perkin Elmer Opera Phenix Plus and Harmony bioinformatics software at UCSC’s Chemical Screening Center. Valid cells were identified as nonborder objects with the presence of nuclear Hoechst staining. Valid regions within objects for quantifying stress granules were defined by those with cytoplasmic staining. G3BP1-positive spots were then defined by filtering for spot width > 0.1 um and relative spot to background intensity > 3. The following was then measured for G3BP1-positive spots: The number of G3BP1-positive spots per cell, size of G3BP1-positive spots, and relative intensity of G3BP1-positive spots to background. N = 4 wells of a 96 well-plate were seeded per genotype and treatment and at least 1,000,000 objects were analyzed per genotype/treatment combination. The resulting values were plotted and statistical analysis was performed on Python.v3.7.7 and Jupyter notebook v6.3.0. Significance was calculated using a Kruskal-Wallis test with Dunn’s multiple comparisons test.

### Cigarette smoke treatment

CSC was obtained from Murty Pharmaceuticals (Cat# nc1560725) at a stock concentration of 40 mg/mL total particulate matter (TPM). HBEC3kts from clone 1 and clone 2 were seeded in a 96 well-plate and grown to 50% confluency. Cells were then treated with 0, 15, 60, and 120ug/mL CSC for three days. Following treatment, the cells were washed twice with DPBS and assayed using CelTiterGlo as described above. Luminescence was normalized to the 0ug/mL control. For each genotype, data was combined from n = 4 of 2 clones, for a total n of 8. Significance was calculated using a Mann-Whitney test using GraphPad Prism.

### RNA-Seq Data Analysis

Raw sequencing reads in fastq files were aligned to a version of the human genome hg38 that has a region of repeats masked to make the *U2AF1* locus alignable^49^, using STAR.v2.7.3a^50^ with the parameters --outSAMtype BAM SortedByCoordinate --twopassMode Basic --quantMode GeneCounts --bamRemoveDuplicatesType UniqueIdentical and the Gencode v33 primary assembly gtf file. Aligned BAM files were indexed with Samtools.v1.10^51^. Mapped reads in BAM files were counted with HTSeq.v0.12.4^52^ for all the annotated genes in gencode.v33.primary_assembly.annotation.gtf with -stranded = reverse and nonunique=none parameters.

### *U2AF1^S34F^* mRNA ratio

Aligned reads from clone 1 and clone 2 HBEC3kts were loaded onto the Integrative Genomics Viewer^53^ (IGV) at the *U2AF1^S34F^* mutational locus. The fraction of A (mutant) nucleotides at this locus obtained from IGV was plotted. Significance was calculated with a Mann-Whitney test on GraphPad Prism.

### Differential expression analysis

Libraries from the same biological replicates of clone 1 samples were sequenced in two different batches, so were read into DESeq2 as technical replicates. Differential expression analysis was performed with DESeq2 v1.40.2^54^ on R v4.3.1^55^ on aligned RNA sequences from clone 1 and clone 2 of our HBEC3kt lines. Gene counts were normalized and a likelihood ratio test calculation was performed to account for batch differences between samples from clone 1 and clone 2. Statistical analysis was performed on expression differences in the following pairwise comparisons: *U2AF1^S34F^ + LacZ* vs. *U2AF1^WT^ + LacZ*, *U2AF1^WT^ + KRAS^G12V^* vs. *U2AF1^WT^ + LacZ*, and *U2AF1^S34F^ + KRAS^G12V^* vs. *U2AF1^WT^ + LacZ*.

### Normalized gene count comparisons

Normalized gene counts for *U2AF1* and *KRAS* were obtained using DESeq2^54^ for each pairwise comparison and plotted with Python.v3.7.7 and Jupyter notebook v6.3.0^56^ . Significance was calculated using Kruskal-Wallis test with Dunn’s multiple comparisons test using Python’s scikit_posthocs module.

### Differential splicing analysis on HBEC3kts

Aligned files from clone 1 technical replicates were merged using samtools merge with default options. .BAM files from all clones were filtered to remove nonstandard chromosomes with samtools v.1.13. Splicing quantification was performed with JuncBASE^57^ v.1.2 beta using the multiprocessing version with the following non-default parameters. First, run_preProcess_by_chr_step1.py was run with --preProcess_options “–unique-j /path/to/intron/coordinates -p 20. Next, disambiguate_junctions.py was run with --by_chr --majority_rules. run_preProcess_step3_by_chr.py was run with --LSF --lsf_queue --min_overhang 6 --num_processes 8 --force --nice. To identify and quantify alternative splicing events in each sample, run_getASEventReadCounts_multiSample.py was run with --sqlite_db_dir/path/to/sqlite_db_dir --jcn_seq_len 240 -p 20 --by_chr. Tables of raw and length-normalized counts of inclusion and exclusion isoforms were created using run_createAS_CountTables.py --jcn_seq_len 240 --num_processes 30. The full JuncBASE commands are in the juncbase_run.sh file provided in the GitHub.

JuncBASE count files were statistically analyzed with the compareSampleSets.py module, using the following non-default commands: --mt_correction BH --which_test t-test --delta_thresh10.0. *U2AF1^WT^+LacZ* samples were compared with the mutant genotypes. Then, redundant splicing events were filtered out using the JuncBASE script makeNonRedundantAS.py. To compare splicing event type distributions between the genotypes, gseasplicing events were filtered for padj < 0.25 and abs(ΔPSI) ≥ 10. We also filtered out junction-only alternative acceptor and alternative donor events, as these events have less read support than other categories. Additionally, we filtered for intron retention events that consisted of known junctions. Statistical differences between splicing event distributions were calculated using a Fisher’s exact test and row-wise Fisher’s exact test with R’s rstatix library.

### GSEA on differentially expressed genes

The log2 fold change (FC) values from each differential gene expression comparison was filtered for adjusted p-value (padj) < 0.05, and the filtered log2 FC values along with gene names were exported as a .RNK file for gene set enrichment analysis using a custom Python script in the GitHub. Gene set enrichment analysis was performed by inputting .RNK files generated from differential expression or splicing analysis into GSEAPreranked on the GSEA v.4.3.2 software^58^. The “Collapse/Remap to gene symbols” option was set to “No_Collapse” and default settings were used for the remaining options. The positive and negative GSEA output tables for each gene set were combined, and the normalized enrichment scores (NES) were filtered for FDR q-value < 0.25 and nominal p-value < 0.05. Filtered NES and the identities of their corresponding gene sets were plotted in heatmaps using Python.v3.7.7 and Jupyter notebook v6.3.0. NA values corresponding to gene sets where NES from certain pairwise comparisons did not pass the filters were replaced with 0 for plotting.

To generate the .RNK file for splicing changes, we took the absolute value of the ΔPSI values produced by compareSampleSets.py and filtered them for padj < 0.25. Unlike differential gene expression, multiple splicing events are possible for a given gene. To convert our results into a.RNK file readable by GSEA, we handled duplicate ΔPSI entries by keeping the entry with the highest abs(ΔPSI) value. The ΔPSI values and gene names were then exported as a .RNK file for GSEA. A list of stress granule RNA and protein genes was obtained from Biancon et al.^21^ and converted to GMX format for GSEA Preranked analysis. The same filtering thresholds were applied to NES scores as in the gene expression GSEA.

### Stress granule protein gene set analysis

A list of stress granule RNA and protein genes was obtained from Biancon et al.^20^. To generate a list of stress granule protein genes reported to also be differentially bound to by *U2AF1^S^*^34^ HBEC3kts, we filtered the stress granule protein gene list for genes that exhibited differential U2AF1 binding from an RNA-IP experiment performed with anti-U2AF1 generated by Palangat et al.^14^. Heatmaps of differentially expressed and spliced genes were then generated with this gene set.

### EIF4A2 isoform visualization

To visualize full-length isoforms of EIF4A2, a GTF of isoforms constructed from FLAIR was obtained from Soulette et al. 2023^17^. This annotation file consists of isoforms constructed from long-read nanopore sequencing of HBECs wild-type and mutant for U2AF1S34F. The GTF was subsetted for entries corresponding to EIF4A2 and converted from hg19 to hg38 coordinates using liftOver. As there were too many isoforms to display, we filtered for the top three isoforms in U2AF1S34F or U2AF1WT cells with the highest normalized read counts.

### U2AF1^S34F^ and KRAS^G12V^ co-occurrence

Lung ADC patient sample mutational statuses was obtained from cBioPortal^59^. Overlapping studies as well as the TSP Nature, 2008 were excluded from analysis. Co-occurrence p value was obtained from cBioPortal’s Mutual Exclusivity analysis and the mutational status of patient samples were plotted with GraphPad Prism.

### *U2AF1^S34F^* and smoking history splicing alteration status

Lung ADC patient sample RNA-seq data was obtained from TCGA and smoking status for the patients was obtained from Campbell et al.^60^. Splicing alteration status was obtained by running JuncBASE to compare lung ADC against matched normal tissues. “jcn_only”, novel intron retention events, and events where more than 25% of the samples were missing data were excluded. Splicing event PSI medians and interquartile range (IQR) were computed from samples with a smoking designation from Campbell et al. A splicing event was considered altered in an individual sample if the PSI – median was more than 1.5xIQR for that event, and the ΔPSI was more than 10% from the median. P-values were calculated using a Mann-Whitney test.

## ACKNOWLEDGMENTS

The short-read Illumina sequencing was carried by the DNA Technologies and Expression Analysis Core at the UC Davis Genome Center, supported by NIH Shared Instrumentation Grant 1S10OD010786-01. Technical support was provided by Dr. Beverley Rabbitts, UCSC Chemical Screening Center RRID SCR_021114. Purchase of the Perkin Elmer Opera Phenix Plus and BioTek EL406 used in this research was made possible through the National Institutes of Health Grant 1S10OD028730-01A1, awarded to the UCSC Chemical Screening Center RRID SCR_021114. Funding for this work came from TRDRP grant T29KT0401 and CRCC grant C23CR5710 to A.N.B. Additional support was provided by NIH/NIGMS R35GM138122 and NIH/NHGRI R25HG006836. S.S.S. is supported by R37CA269754 and I.J.F was supported by 5T32GM133391.

We would like to thank Drs. Christina M. Towers, Judith Campisi, and Birgit Schilling for providing critical advice and feedback on early stages of this study.

## SUPPLEMENTAL INFORMATION

**Document S1**. Figures S1–S3.

**Table S1**. Excel file containing data for figures plotted with GraphPad Prism. Related to Figures 1A, 1D, 2D, 3A, 3D, 4B; Figures S2 F, S3 C-E.

**Table S2**. Excel file of Log2 fold change values for HBEC3kt lines from DESeq2. Related to Figure 1C, Figure S3 A.

**Table S3**. Excel file containing fluorescent imaging measurements. Related to Figures 3B-C.

## AUTHOR CONTRIBUTIONS

E.H.R., C.E.L., and A.B. performed experiments. C.E.L., E.H.R., S.M., A.B.,C.F., C.M.S., and A.M.T. analyzed data. A.N.B. supervised this study. E.H.R. and A.N.B. conceived of this study. C.E.L. wrote and edited original draft. C.E.L., E.H.R., and I.J.F. wrote methods. I.J.F performed the *in vivo* xenografts studies under supervision of S.S.S.. I.J.F, S.S.S, E.H.R.,C.F., and A.N.B. provided critical feedback on the manuscript.

## DECLARATION OF GENERATIVE AI AND AI-ASSISTED TECHNOLOGIES IN THE WRITING PROCESS

During the preparation of this work the author(s) used Gemini (Google) in order to check for grammar and improve text. After using this tool/service, the author(s) reviewed and edited the content as needed and take(s) full responsibility for the content of the publication.

---

**S1.**
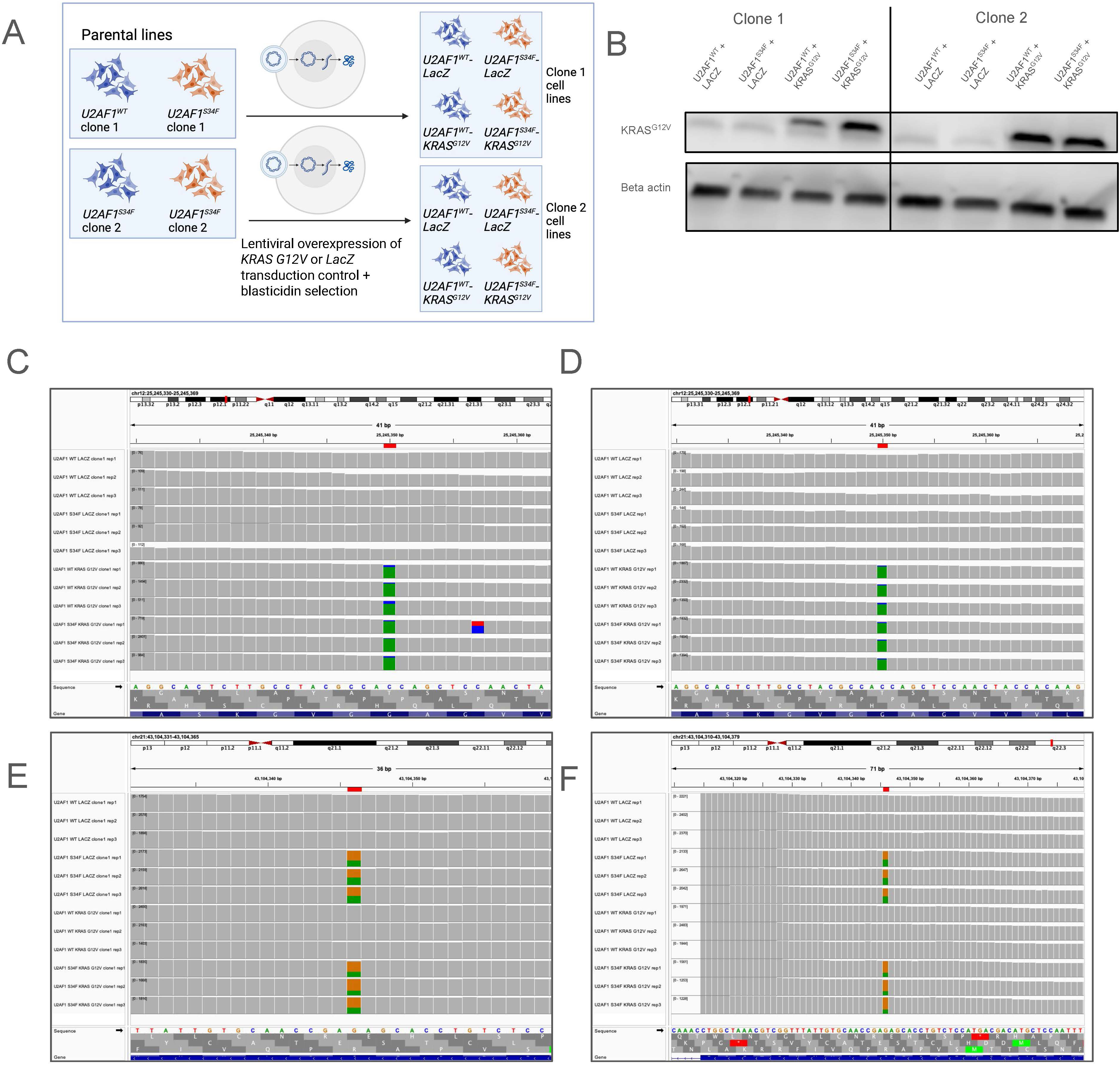
Validation of cell lines for *U2AF1^S34F^* and *KRAS^G12V^*. **(A)** Schematic of cell line generation from 2 parental clones of *U2AF1^S34F^*and *U2AF1^WT^* HBEC3kts. (**B)** Western blot validation with *KRAS^G12V^* -specfici antibody in clone 1 and 2 HBEC3kts. **(C)** IGV validation of *KRAS^G12V^* mutation for clone 1 HBEC3kts. **(D)** IGV validation of *KRAS^G12V^* mutation for clone 2 HBEC3kts. **(E)** IGV validation of *U2AF1^S34F^* mutation for clone 1 HBEC3kts. **(F)** IGV validation of *U2AF1^S34F^* mutation for clone 2 HBEC3kts. The loci of *U2AF1^S34F^* and *KRAS^G12V^*mutations are highlighted with a red bar at the top of the IGV browser shot. Colored bars in the coverage tracks denote the presence of deviations from the reference sequence.

**S2.**
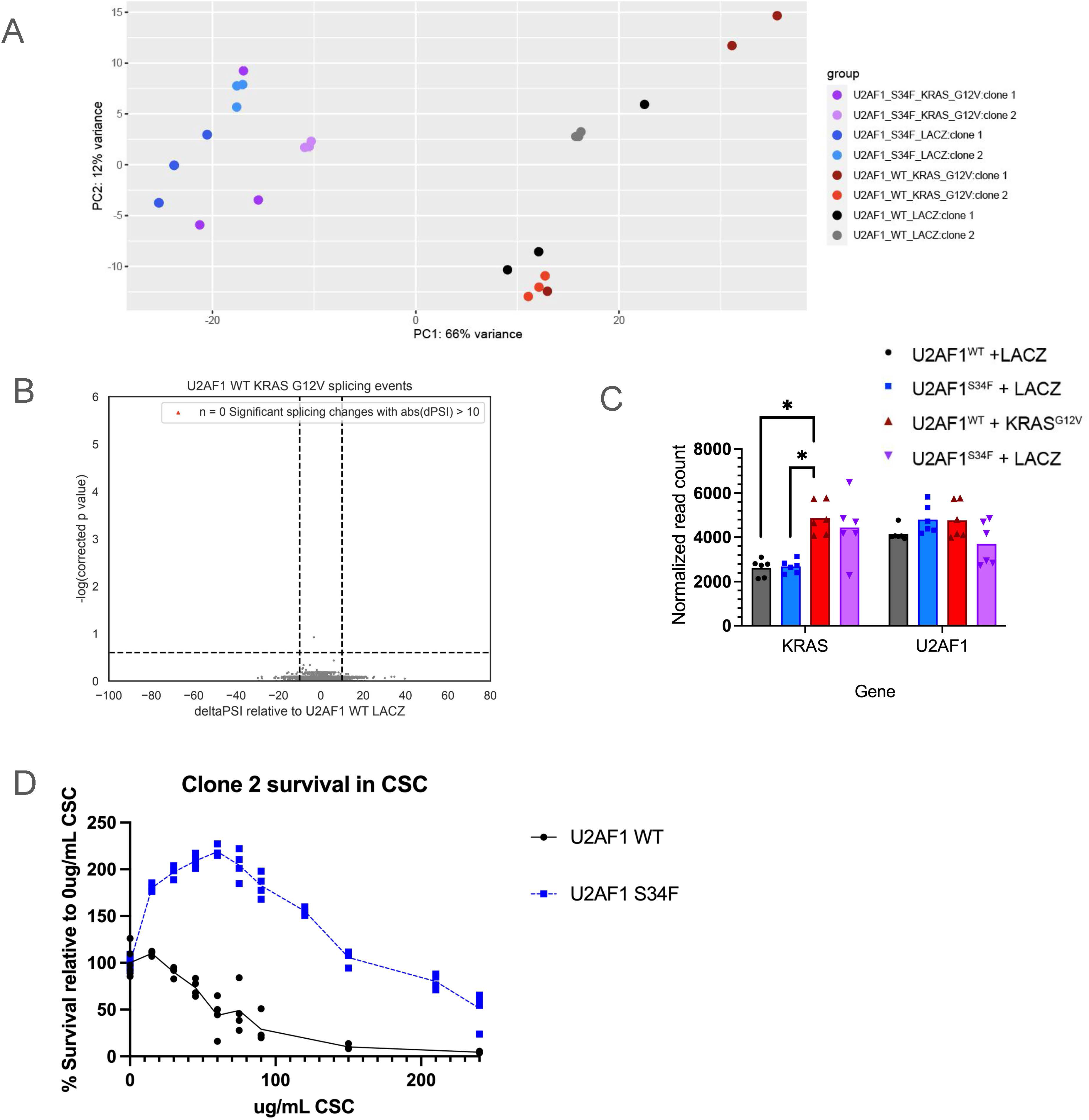
Gene expression and viability differences in HBECs with *U2AF1^S34F^*and *KRAS^G12V^*. **A,** PCA of clone 1 and clone 2 gene expression. (B) Volcano plot of splicing events detected by JuncBASE for U2AF1 WT KRAS G12V HBEC3kts. (**C)** Normalized gene counts of *KRAS* and *U2AF1* in HBEC3kts.KW + Dunn’s multiple comparisons test. **(D)** Survival of clone 2 HBECs wild-type and mutant for U2AF1 under three days of exposure of 0 to 240ug/mL CSC. Percent survival is measured by obtaining the percent viability of each time point relative to 0ug/mL CSC. * P ≤ 0.05

**S3.**
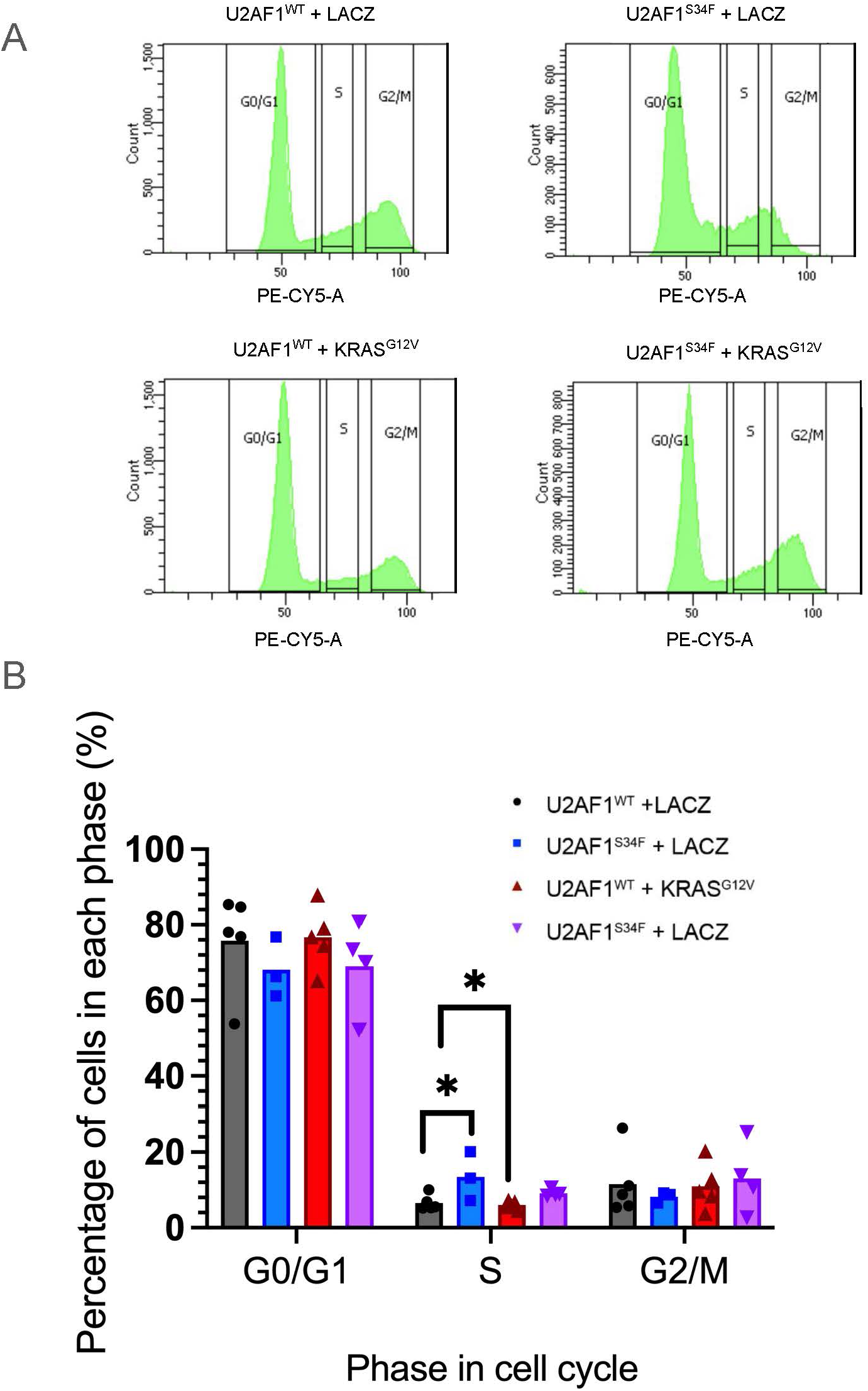
Cell cycle differences in *U2AF1^S34F^* and *KRAS^G12V^*HBECs. (**A)** Counts of the number of events in each phase of the cell cycle from propidium iodide (PI) staining and flow cytometry for each HBEC genotype. **(B)** Percentage of events from A in each of the cell cycle phases for HBECs. KW + Dunn’s multiple comparisons test. * P ≤ 0.05,

**S4.**
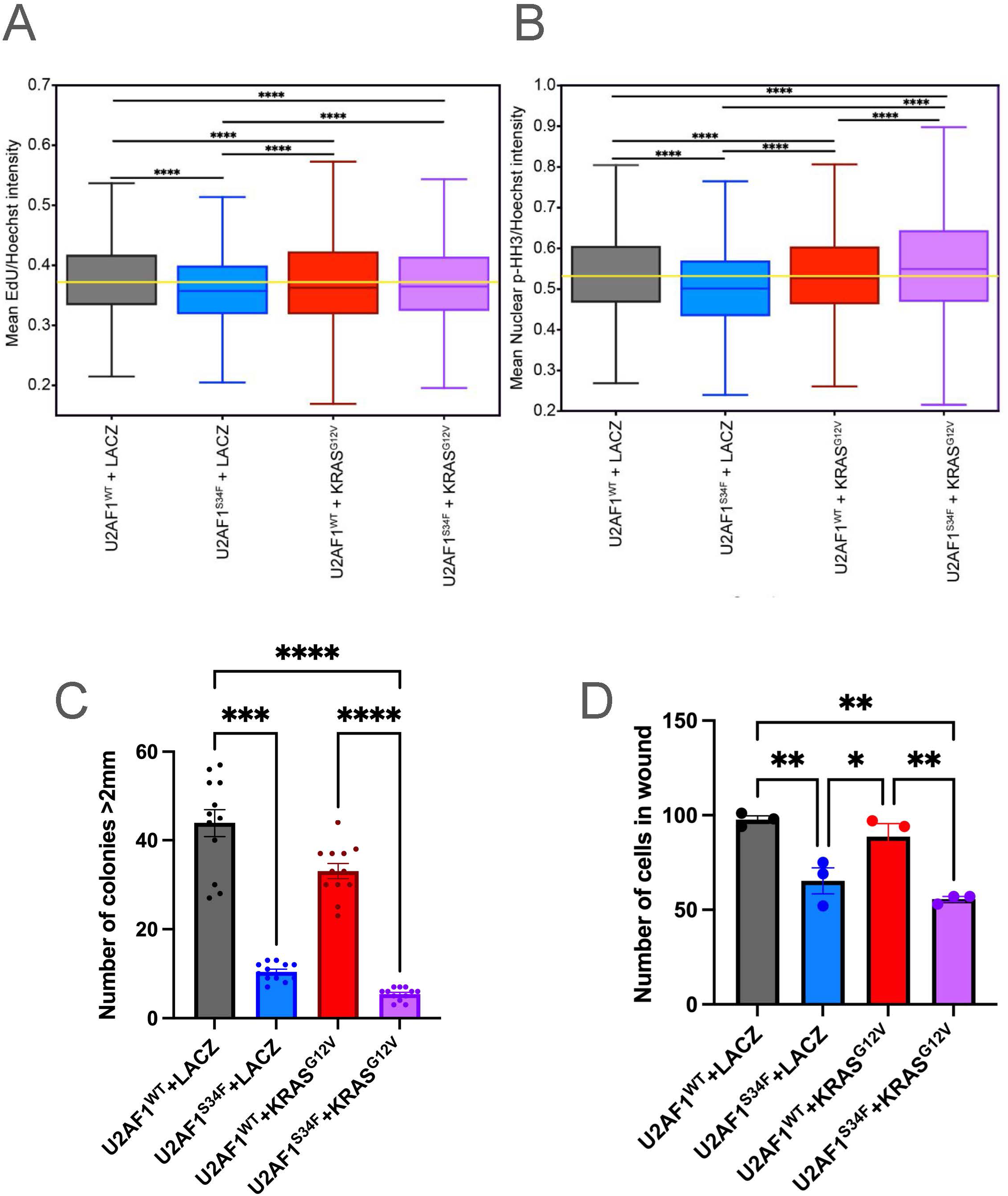
Additional assays of HBEC3kts with *U2AF1^S34F^* and *KRAS^G12V^*mutation. (**A)** Mean EdU intensity of cells, normalized by cell density. (**B)** Mean PHH3 intensity, normalized by cell density. Yellow lines correspond to the median of the wild-type control. The middle line in the body of each boxplot represent medians of each genotype, box limits represent quartiles, and whiskers represent the range of the most extreme, non-outlier data points. **(C)** Clonogenicity assay of clone 2 HBEC3kts. Bars represent average numbers of colonies larger than 2mm. **(D)** Potential of clone 2 HBEC3kts to invade wound after 3 hours in 2-D culture. Bars represent average number of cells in wound. Error bars represent SEM. * P ≤ 0.05, ** P ≤ 0.01, *** P ≤ 0.001, **** P ≤ 0.0001. See also Table S1, Table S2. A-B: Kruskal-Wallis with Dunn’s multiple comparison. C-D: one-way ANOVA with Holm-Šídák’s multiple comparisons test

